# Profiling copy number alterations in cell-free tumour DNA using a single-reference

**DOI:** 10.1101/290171

**Authors:** Alan J Robertson, Qinying Xu, Sarah Song, Devika Ganesamoorthy, Derek Benson, Wenhan Chen, Kaltin Ferguson, Katia Nones, Sunil R Lakhani, Peter T Simpson, Nicola Waddell, John V Pearson, Lachlan J.M. Coin

## Abstract

**Background:** The accurate detection of copy number alterations from the analysis of circulating cell free tumour DNA (ctDNA) in blood is essential to realising the potential of liquid biopsies. However, currently available approaches require a large number of plasma samples from healthy individuals, sequenced using the same platform and protocols to act as a reference panel. Obtaining this reference panel can be challenging, prohibitively expensive and limits the ability to migrate to improved sequencing platforms and improved protocols.

**Methods:** We developed qCNV and sCNA-seq, two distinct tools that together provide a new approach for profiling somatic copy number alterations (sCNA) through the analysis of cell free DNA (cfDNA) without a reference panel. Our approach was designed to identify sCNA from cfDNA through the analysis of a single plasma sample and a matched normal DNA sample -both of which can be obtained from the same blood draw. qCNV is an efficient method for extracting read-depth from BAM files and sCNA-seq is a method that uses a probabilistic model of read depth to infer the copy number segmentation of the tumour. We compared the results from our pipeline to the established copy number profile of a cell-line, as well as the results from the plasma-Seq analysis of cfDNA-like mixtures and real, clinical data-sets.

**Results:** With a single, unmatched, germline reference sample, our pipeline recapitulated the known copy number profile of a cell-line and demonstrated similar results to those obtained from plasma-Seq. With less than 1X genome coverage, our approach identified clinically relevant sCNA in samples with as little as 20 % tumour DNA. When applied to plasma samples from cancer patients, our pipeline identified clinically significant mutations.

**Conclusions:** These results show it is possible to identify therapeutically-relevant copy number mutations from plasma samples without the need to generate a reference panel from a large number of healthy individuals. Together with the range of sequencing platforms supported by our qCNV+sCNA-Seq pipeline, as well as the Galaxy implementation of this solution, this pipeline makes cfDNA profiling more accessible and makes it easier to identify sCNA from the plasma of cancer patients.

## Introduction

Somatic copy number alterations (sCNA) are an important class of mutation in cancer [1, 2]. Specific sCNA such as the amplification of *HER2* in breast cancer or the Androgen Receptor (*AR*) in prostate cancer have both been linked to better outcomes from specific therapies and increased five-year survival rates [3-6]. Determining the specific copy number mutations present in an individual’s cancer is essential for fully understanding each patient’s disease.

Characterisation of the cell-free DNA (cfDNA) in the bloodstream of cancer patients can offer insights into the specific molecular events associated with each patient’s disease without the need for invasive surgeries [7-11]. Analysis of the circulating tumour DNA (ctDNA) component of cfDNA, can make it possible to determine an individual’s response to specific treatments [12], and to monitor the evolution of the patient’s disease [13, 14]. While the importance of sCNA are well established, much of the work surrounding the use cfDNA to characterise a patient’s disease has focused on the analysis of somatic single nucleotide variations (SNV) [5, 15-17].

Several methods have identified copy number changes from cfDNA by comparing the DNA from the patient’s plasma to a panel of reference samples collected from the plasma of groups of healthy individuals, that have been processed and characterised using the same technology[8, 10, 16, 18]. These methods identify sCNA segments by determining regions in the cfDNA that are significantly different from the reference panel (Z-score). This Z-score based approach has been effectively used across a range of different cytogenetic settings and is routinely used in prenatal screening [18, 19].

The most widely used method for determining sCNA from the cfDNA of cancer patients using high throughput sequencing, is the Z-score based approach implemented in plasma-Seq [8, 14, 20, 21]. plasma-Seq can identify genome-wide sCNA from ultra-low cfDNA sequence coverage (0.05x – 0.2x) cfDNA data-sets [8, 21, 22]. Analysis of a cell-line processed to resemble cfDNA (DNA samples fragmented to 150 -250 bp) with plasma-Seq, demonstrated that it was able to recapitulate the established copy number profile of the cell-line across a range of samples that contained varying amounts of tumour DNA [8]. Together, these attributes have made plasma-Seq an effective and popular approach for identifying sCNA in cfDNA, and have allowed researchers to characterise the copy number profiles from the plasma of patients suffering from a range of different cancer types [8, 10, 14, 20, 21].

However, the generation of a panel of library-matched, technology-matched, and sex-matched data-sets from a group of healthy individuals can be prohibitively difficult for smaller laboratories to source and generate. Moreover, migration to a new sequencing technology, alterations to sequencing chemistry, or improvements to the processes for manipulating cell-free DNA mandates the periodic generation of a new reference panel. Moreover, this reference panel based approach can make it difficult to separate germline copy number variants (CNV) from somatic sCNA.

In this manuscript, we describe the qCNV+sCNA-seq pipeline, which identifies sCNA without requiring a reference panel of healthy plasma samples. This approach identifies sCNA from the cfDNA of a cancer patient relative to a single reference sample, such as the cfDNA sample from a healthy, unrelated individual, or a sample of normal germline DNA from the same patient. We show this pipeline can characterise sCNA through the analysis of artificial, cfDNA-like mixtures and clinical cfDNA samples. We also describe a web-based galaxy implementation the qCNV+sCNA-Seq pipeline [23], which can be used by labs without dedicated bioinformatics resources. Together the versatility of this approach makes it possible for more researchers to adopt cfDNA sequencing to identify sCNA.

## Methods

We developed a new pipeline composed of two novel tools qCNV and sCNA-seq (Figure 1). qCNV is a memory and time efficient method for quickly determining the number of reads that align to specific regions of the genome; while sCNA-seq takes the output from qCNV and uses the distribution of reads from the ‘normal’ reference sample to identify sCNA from the cfDNA sample. Both tools operate independently from one another, allowing researchers to make use of the advantages each tool offers for their own pipelines. The counts file from qCNV can be used for any purpose, and sCNA-Seq can use the correctly formatted counts file from an alternative program, but together these tools offer a fast pipeline able to detect sCNA from low coverage cfDNA samples.

**Figure 1.**
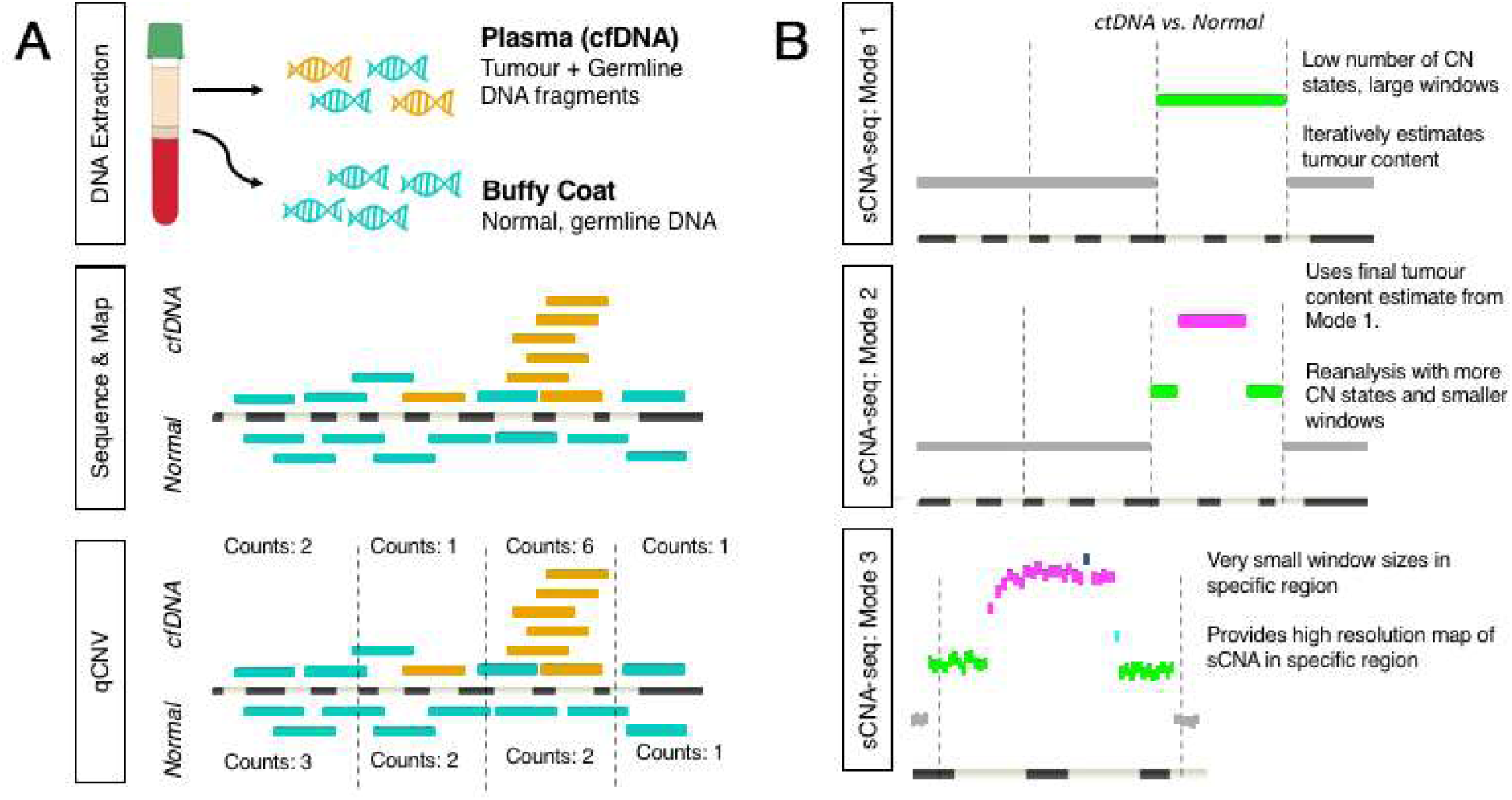
The qCNV+sCNA-Seq Workflow. The pipeline is made up of two separate tools and three distinct stages. (A) The first stage uses qCNV to determine the number of reads that align to a 1000bp window in both the cfDNA and normal germline samples. (B) The second stage takes the output from qCNV and passes it through the whole genome training mode of the second tool, sCNA-Seq. This mode uses larger window sizes and a smaller number of copy number states to better determine the proportion of tumour content in a sample. The third stage uses the tumour content estimation to run sCNA-Seq with a smaller window size (down to 2 kb) and a higher number of potential copy number states to provide a high resolution annotation of the tumour genome through the analysis of the cfDNA. This can be run across the entire genome (Mode 2) or for a specific region of the genome (Mode 3).

### Data from previous plasma-Seq experiments

Heitzer et al [8] generated a series of plasma-like DNA samples to determine the capacity of the plasma-Seq approach to characterise cfDNA samples that contained varying amounts of tumour derived DNA (Additional File 1, Supplementary Table 1). These mixtures were generated by mixing DNA from the HT-29 tumour cell-line with DNA from an unmatched normal cell line at varying proportions and subsequently mechanically shearing the DNA in order to generate fragment lengths similar to those observed in cell-free DNA (150 -250 bp). These samples were then sequenced with the MiSeq platform (1x 150bp). We downloaded this data from the European Genome-Phenome Archive (accession: EGAD00001000364) [24]. We also downloaded raw sequence data from 11 cfDNA samples collected from 7 prostate cancer patients, and 9 cfDNA samples from healthy females using the same accession. These datasets were mapped to the HG19 annotation of the human genome with BWA, using the internal pipeline detailed below. The coverage of these samples ranged from 0.06X -0.29X coverage (Additional File 1, Supplementary Table 1).

### Matched exome data from tumour, metastasis, cfDNA and germline samples

Butler et. al [17] generated exome sequencing data on primary tumour, metastatic, cfDNA and germline samples from two patients (Additional File 1, Supplementary Table 1). This data set contained data from metastatic breast cancer and sarcoma. Mapped bams were downloaded from the European Nucleotide Archive [25] (accession PRJEB8969). The mapping and processing of these files were described in the original publication [17], the coverage of these samples ranged from 118x to 309x.

### Visualization of the established copy number profile for the HT-29 cell-line

An independent annotation of the complete copy number profile of the HT-29 cell-line was downloaded from the COSMIC Cell Lines Project (CCLP) [26], using the Sample ID: COSS905939. This whole genome annotation of the sCNA in HT-29 was generated the CCLP through a PICNIC analysis [27] of microarray data produced by the Affymetrix SNP6.0 array. Unlike the results from plasma-Seq or qCNV+sCNA-seq, which analysed short degraded fragments of DNA in the plasma, the DNA used here was, genomic collected from intact cells, in a pure cell-line. The copy number profile generated by the CCLP represents the gold standard copy number for the HT-29 cell-line. To visually represent this information, the data file containing the absolute copy number of each segmented was plotted with ggplot2 [28]. To allow readers to fairly compare the results from different platforms each chromosome was condensed to the same width as the corresponding chromosome in the sCNA-seq figure.

### Generation of a sex matched, germline reference, used in the comparison to EGAD00001000364

As the EGAD00001000364 cohort did not contain any normal, germline samples, we made use of an existing ultra-low coverage, sex-matched sample of germline DNA that our group had previously sequenced with the MiSeq platform. This sample was collected as part of a study to follow-up patients diagnosed with breast cancer. The collection of breast cancer samples, by the Brisbane Breast Bank, was approved by the Human Research Ethics Committee at the University of Queensland (approval number: 2005000785). A sample of the patient’s germline DNA was collected from a buffy coat sample and was used as our unmatched, normal reference. The DNA from this sample was extracted with Qiagen’s DNeasy Blood and Tissue kit and was sequenced using the second iteration of the MiSeq sequencing chemistry. These raw fastq files were then passed through the same mapping pipeline as the raw plasma-Seq data from EGAD0000100036. While low coverage (0.92X), our reference sample was still considerably deeper than any of the plasma-Seq samples (Additional Supplementary Table 1). To achieve a comparable depth this reference sample was down-sampled using Samtools [29]. The final coverage of the downsampled reference sample used in these comparisons was 0.18X.

### Sequence Alignment and Post-Processing

All raw fastq files, (both public and sequenced in this study) were mapped to the genome (HG19) with BWA-MEM [30]. The resulting SAM file produced by BWA was sorted and converted into a bam file using Samtools [29]. Samtools was used to filter out supplementary alignments and index the filtered bam. The duplicate reads in this filtered bam were then identified using the MarkDuplicates function of Picard [31] The filtered, sorted, duplicate marked bam file was then characterised by qCoverage and qProfiler [32] to determine the quality and coverage of the mapped samples. When required, down-sampling of the final bams, was performed with the view -s function of Samtools.

### Detection of sCNA with the qCNV+sCNA-seq pipeline

This pipeline was developed to quickly analyse pairs of cfDNA/normal bams and identify sCNA. The pipeline is made up of qCNV and sCNA-seq. These tools were designed to complement each other, however both applications can be run independently of each other.

### qCNV

qCNV is an efficient java tool designed to determine the number of reads that align to specific regions in the genome. This method divides whole genome into windows of a fixed size and counts the number of reads that begin in each window. The size of the windows is defined by the user at the beginning to the experiment. The window size used in each of these experiments is 1000 bp.

### sCNA-seq

sCNA-seq characterizes the copy number profile of cfDNA sequencing data through the analysis of raw read count produced by qCNV. This method allows for the detection of both small scale and larger mutations. In our model, we used the normalized read counts from user-defined windows, to estimate the proportion of tumour DNA in the cfDNA sample and in cfDNA/normal pairs from the same individual, distinguish sCNA from CNVs. sCNA-seq utilises the hidden Markov model from cnvHitSeq [33], although the read-depth modelling is substantially different.

### Expectation-maximization (EM) model for identifying sCNA from circulating tumour DNA in cfDNA samples

We divide each chromosome into windows of a fixed size (described further below). We consider the observed data to consist of the number of reads which begin in each window in normal and plasma samples, N_i_ and T_i_ respectively. Define N = ∑ N_i_ and T = ∑ T_i_ as the total read counts for normal and plasma samples. The hidden states in our model consist of the ratio of the tumour copy number to the normal copy number in each window rcn_j_ = {0,0.5,1,1.5,2,2.5,..}, so that rcn = 1 corresponds to no amplification or deletion. We also define two parameters in our model: the tumour *purity*, which is the proportion of cell-free DNA which is derived from tumour, and the ploidy *ratio* as the average relative copy number state across the genome (which can be considered as the ratio of observed ploidy to normal ploidy of 2).

We model the expected number of reads in the plasma sample in each window using a beta-binomial distribution

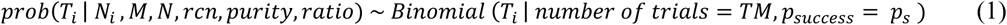

where

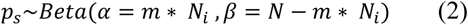

where we have defined a multiplication factor m as

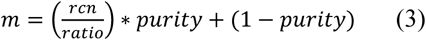

This multiplication factor can be thought of as a way to calculate the expected proportion reads in a given window in the plasma genome, conditional on the proportion of reads observed in the same window in the normal sample. The first term, (rcn/ratio)*purity, reflects that a greater proportion of reads are expected to come from the tumour portion in proportion to the increase in copy number relative to the average. The second term (1-purity) in this equation is the contribution to this proportion from the non-tumour derived component of plasma.

The beta distribution models our uncertainty in the estimate of the probability of selecting a read in a given window in the normal genome. This uncertainty scales with the number of independent reads. We introduce a parameter to adjust for any non-independence (e.g. caused by PCR), called Beta-down weight (BDW), such that

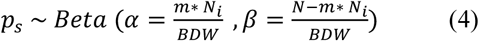

One example of use of this parameter is to adjust for counting both ends of a paired end read (in this case BDW = 2). However, we have observed that BDW is also useful for modelling over-dispersion observed in both whole genome and exome-capture datasets.

We adopt the hidden Markov model described in Bellos et. Al [33]. The emission probabilities are defined by equation 1. This model allows transitions between different copy number states according to a globally defined transition rate matrix, such that a per-base transition rate is defined, and the transition probability matrix is defined as a matrix exponential of the rate matrix scaled by the distance measured between the mid-point of each window.

The ratio is initialized to 1, and the purity is initialized to 0.99. We use the forward-backward algorithm to calculate the posterior probability of using each relative copy number state at each window, w_i_ (rcn_j_), which we use as a weight to assign each data point (T _i_, N _i_) to each copy number state. We then use a gradient descent algorithm in order to calculate the tumour purity and ratio which maximize the probability of this data, i.e. the combination of purity and ratio which maximizes

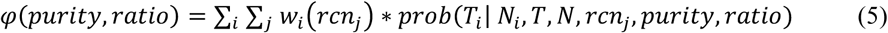

We iterate these two steps until convergence of the purity and ratio estimates (such that the difference between iterations is less than 0.01), and value of the objective function or until a pre-defined maximum number of iterations is reached.

### Analysis of cfDNA sCNA-seq modes

sCNA-seq can be run in three modes (Supplementary Methods). The first mode is a training mode which estimates the proportion and ratio of tumour DNA in the sample and provides a low resolution sCNA segmentation genome-wide. The second mode uses this information to characterise the tumour genome at a higher resolution and with an increased number of copy number states (Figure 1). The third mode allows the user to perform a high-resolution analysis of a specific section of the genome. Like the second mode, this high-resolution survey mode, requires tumour content information produced in the whole genome training mode.

### GC correction

To address for the impact of GC biases on the analysis on the low coverage samples that compared cfDNA to the genomic DNA from intact cells, a form of GC correction was developed to apply to the results from qCNV. This correction used a polynomial regression (with a quadratic model), to adjust the results for the amount of GC content for each of the 1000 bp regions of the genome in both the cfDNA and reference samples. As the effects of GC bias differ between runs and batches, this correction should be run under the user’s discretion. For all samples in the EGAD00001000364 data-set, GC correction was applied, however it was not needed in the samples described by Bulter et. al [17]

### Quality Control

In order to assess the quality of the results from qCNV + sCNA-Seq analysis of whole genomes, we introduced a quality control measure to account for excessive copy number switching. This can occur if the model is unable to resolve the correct tumour purity as a result of unmodeled variation in the data. As an example, we observed this behaviour when modelling publicly available exome data [17] for which the plasma and matched normal were assayed on different versions of an exome capture array. We flag the copy number segmentation as unreliable if the observed number of distinct sCNA exceed 5% of the number of potentially distinct sCNA (which is just the total number of windows used in the sCNA-Seq analysis).

### Comparing copy number profiles

Determining the exact level of overlap between two different profiling methods (the official CCLP PICNIC copy number profile for the HT-29 cell-line [26], the pre-existing plasma-Seq results [8] and the results from qCNV+sCNA-seq) was done at a base pair resolution and involved comparing the exact position of all of the sCNA identified in each analysis to the total sCNA identified in the corresponding analysis. In this comparison, sCNA were defined as either a gain or a loss, with both copy number mutations representing a genomic region that did not contain the same copy number state as the defined base-ploidy of the sample. The results from the qCNV + sCNA-Seq pipeline were typically designated as the reference for consistency. The number of overlapping bases between the reference sCNA and those identified in one of the other analyses were used to measure sensitivity and specificity.

The results from original plasma-Seq analysis [8] and the independent copy number profile from the CCLP analysis of the HT-29 cell-line [26] had to be reformatted to allow for this comparison. For the sCNA identified by the CCLP, this involved defining any segment lower than the base ploidy as a loss, while segments with a copy number estimate greater than the base ploidy were classed as gains. As the results from plasma-Seq do not contain any estimates of absolute copy number changes, a loss was defined as region with a segmental Z-score of < 5.00, while gains had a Z-score of >5.00, a method previously used when comparing the copy number results from plasma-Seq to another technology [34]. In both samples, any reads aligning to the Y chromosome were removed. The X chromosome was retained due to the importance of this chromosome in prostate cancer.

### Galaxy Implementation

The java applications used for analysis have been wrapped for use by the Galaxy scientific workflow system and added to a Galaxy server (http://www.genomicsresearch.org/galaxy) under the menu item “Plasma CNV Pipeline” [23]. The values required to run these programs have been pre-set, likewise the small amounts of preprocessing to allow these tools to seamlessly communicate with one another have been put in place. This Galaxy server was launched as part of a cluster on the NeCTAR research cloud using the GVL(1) launch process.

## Results

### The sCNA-Seq recapitulates the established copy number profile of a plasma-like cell-line using a single plasma reference sample

We first assessed whether a single reference sample of cfDNA from an unrelated, healthy individual was sufficient to identify known sCNA from a cancer cell-line with an established copy number profile. To achieve this, the qCNV + sCNA-Seq pipeline was used to analyse DNA from the triploid HT-29 cancer cell-line that had been fragmented to resemble the plasma cfDNA, albeit at 100% tumour purity [8], using a single sample from the plasma-Seq reference panel as a reference. Our single reference approach recapitulated the same copy number profile present in the official COSMIC Cell Lines Project (CCLP) analysis of this cell-line, as well as the corresponding plasma-Seq annotation of the same sample (Figure 2 A,B,C). Using a 150 kb window size, the sCNA-seq pipeline identified clinically significant sCNA known to be present in HT-29, such as the gain of chromosome 11, as well as the amplification of 8q24 -the region containing the *MYC* oncogene [35, 36].

**Figure 2.**
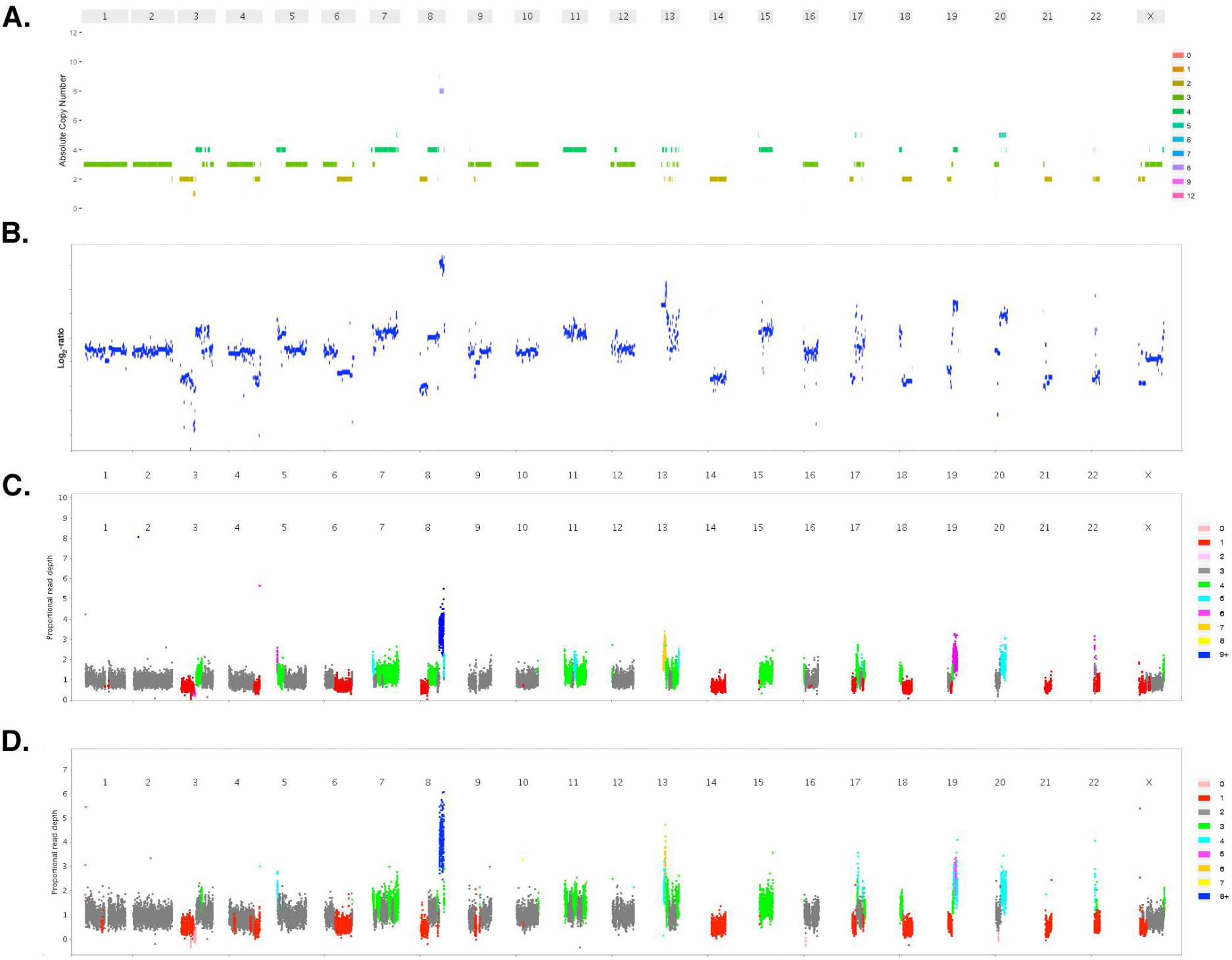
The copy number profile of HT29 cell-line by microarray, plasma-Seq and qCNV+sCNA-seq. In each figure the X-axis represents each chromosome in the genome. To allow for a direct comparison, the spacing of the results from plasma-Seq and the CCLP have been adjusted to match the results from qCNV+sCNA-seq. (A) Copy number profile of the HT-29 cell-line as determined by the CCLP. The Y-axis and the colour of each segment represents the absolute copy number of that region (B) The copy number profile from the plasma-Seq analysis of the pure cell-line and the panel of normal plasma samples. This image was originally published as additional file 3 in plasma-Seq publication [8]. The Y-axis shows the log2-ratio for each segment (C) The copy number profile from the qCNV+sCNA-seq pipeline of the plasma-like cell-line and against a single cfDNA sample from a healthy individual (a member from the plasma-Seq reference cohort), assuming a triploid model. The Y axis and segment colour shows the absolute copy number for each segment (D) The copy number profile from the qCNV+sCNA-seq analysis of the plasma-like cell-line and a single reference sample, prepared from intact normal cells. As information about the ploidy of a tumour is not always known, this sample was analysed as a diploid. Despite incorrectly assigning the ploidy, we were still able to capture the same broad copy number profile. In all the figures produced by sCNA-Seq, the colour of the segment represents a distinct copy number state.

When compared to the sCNA identified in the PICNIC analysis of the high quality, genomic, cell-line DNA used in the CCLP analysis (Figure 2A) [26], sCNA-seq obtained an 87% base-pair sensitivity and 87% base-pair specificity (Table 1). Moreover, both annotations classified similar amounts of the genome as being altered by a copy number mutation (CCLP -1323.5 Mb, sCNA-Seq -1334.7 Mb), despite the fact sCNA-Seq identifying fewer sCNA overall. In contrast, the published plasma-Seq results, which used a reference panel of plasma samples from 9 healthy women classified a higher proportion of the genome as being altered by some form of copy number mutation (1727.3 Mb). When compared to the sCNA identified by the CCLP, the plasma-Seq results had a lower base-pair specificity (74%) but a slightly higher (93%) base-pair sensitivity (Table 1, Figure 2B), suggesting that in this experiment, the model sCNA-seq was able to provide a more accurate representation of the HT-29 cell-line.

**Table 1.** The HT-29 Copy Number Profile as determined by microarray, plasma-Seq and sCNA-seq.

|  | No of.<br>sCNA | sCNA<br>Length | Gain Length | Loss Length | Overlap CCLP<br>(sens/spec) |
| --- | --- | --- | --- | --- | --- |
| **CCLP**<br>**(Genomic DNA)** | 249 | 1323.5 Mb | 728.9 Mb | 594.6 Mb | 100/100* |
| **plasma-Seq**<br>**(cfDNA reference)** | 429 | 1727.3 Mb | 924.9 Mb | 802.3 Mb | 74/93 |
| **qCNV + sCNA-Seq**<br>**(cfDNA reference, ploidy =3)** | 52 | 1334.7 Mb | 732.7 Mb | 602.0 Mb | 87/87 |
| **qCNV + sCNA-Seq**<br>**(Germline reference, ploidy = 3)** | 78 | 1435.4 Mb | 848.2 Mb | 587.4 Mb | 89/82 |
| **qCNV + sCNA-Seq**<br>**(Germline reference, ploidy = 2)** | 80 | 1314.0 Mb | 730.5 Mb | 583.5 Mb | 82/82 |

To ensure that this result was not the consequence of using an exceptional sample as our single reference, we repeated this analysis using each of the remaining 8 ultra-low coverage (0.13 – 0.29X) healthy plasma samples as a single-reference. The copy number profiles from each of these 9 analyses are highly concordant (Supplementary Figure 1, Supplementary Table 2). When compared to the copy number profile from the CCLP, each of these comparisons confidently reflected the established copy number profile of this cell-line (Sensitivities: 85-92, Specificities 85-95) (Supplementary Table 2). Likewise, each of these analyses revealed a similar proportion of the genome altered by sCNA (1278.0 Mb -1378.3 Mb). Together these results show that comparing plasma-like tumour DNA to the cfDNA from a single, healthy individual can identify known copy number mutations in ultralow coverage samples.

### sCNA-seq determines the known sCNA of a plasma-like cell-line using a white blood cell DNA reference

Having shown the feasibility of using a single, unrelated plasma sample as a reference sample, we next investigated the feasibility of using DNA obtained from intact white blood cells as a reference. The experimental advantage of this approach is that it allows normal DNA from white blood cells to be extracted at the same time as cell-free DNA from the same blood draw, making it possible to compare matched pairs of cfDNA and germline DNA. As the samples used to benchmark plasma-Seq did not contain any ‘normal’ germline DNA sample, we used a sex-matched, low coverage WGS sample of normal DNA that had been generated in-house using the same sequencing platform. We down-sampled this file (0.18X) to match the coverage obtained for the reference plasma samples in order to not produce biased results due to greater matched normal read depth.

Using this low coverage, unmatched normal DNA sample as a reference, sCNA-seq obtained an 89% base-pair sensitivity and 82% specificity when compared to the CCLP copy number profile (Figure 2C, Table 1), and when the sample was analysed as a triploid. As ploidy of a tumour may not be known a-priori, we re-analysed this sample using the default diploid model, which led to a slightly reduced sensitivity of 82%. The copy number profile generated from the triploid and diploid models were very similar (Figure 2 C,D). These results suggest that sCNA can be identified from the analysis of the cfDNA in the plasma when compared to intact normal germline DNA, and that it is robust to the mis-specification of tumour ploidy.

### sCNA-seq identifies known copy number mutations across a dilution of cfDNA-like samples

The proportion of tumour derived DNA in cfDNA is highly dynamic. In some patients 90% of the cfDNA can be tumour derived, while in other cancer patients these fragments can almost be undetectable [11, 16, 17, 37, 38]. Having shown the accuracy of the method with a pure ctDNA-like sample (100% cell-line DNA mixture), we next considered the performance of our approach at lower concentrations of plasma-like DNA. To achieve this, we analysed the same ultra-low coverage WGS mixtures (0.11X-0.17X) used to benchmark plasma-Seq with the qCNV+sCNA-seq pipeline, and using our sex-matched, germline reference sample (Figure 3). As with the results from plasma-Seq, as the proportion of tumour cfDNA decreased, we detected a smaller number of copy number mutations, and a smaller proportion of genome altered by these events. The 50% ctDNA mixture was found to represent the pure ctDNA sample with 68% specificity, dropping to 18% in the 5% ctDNA mixture (Table 2). The sensitivity of the approach remained high across the mixtures, only dropping to 94% in the 10% ctDNA mixture and 76% for the 5% ctDNA mixture. A similar pattern was observed for the plasma-Seq analysis, with sensitivity falling to 15% for the 5% mixture, but specificity remaining above 90%. Both plasma-Seq and sCNA-seq suffer from a degree of over-prediction when there is no or very little ctDNA in the mixture, although in this case qCNV+sCNAseq identifies a much smaller amount of spurious sCNA at these very low levels of tumour purity (Table 2). Some of these predicted sCNA at very low levels of tumour purity may be due to germline differences between the patient and unrelated reference.

**Table 2.** The capacity of plasma-Seq and qCNV+sCNA-seq to resolve the copy number profile of cell-free like DNA across a range of mixtures.

|  | plasma-Seq<br>Number of events | plasma-Seq<br>Length of events | plasma-Seq<br>Self consistency*<br>Sensitivity / Specificity | qCNV+sCNA-seq<br>Number of events | qCNV+sCNA-seq<br>Length of events | qCNV+sCNA-seq<br>Self consistency*<br>Sensitivity / Specificity |
| --- | --- | --- | --- | --- | --- | --- |
| **100% HT-29 Cell-line DNA** | **429** | **1727.3Mb** | **100/100\*** | **80** | **1314.0 Mb** | **100/100\*** |
| **50% Cell-line DNA** | N/A** | N/A** | N/A** | 34 | **935.6 Mb** | 68/97 |
| **20% Cell-line DNA** | **69** | **775.7Mb** | 38/95 | 25 | **328.5 Mb** | 25/95 |
| **10% Cell-line DNA** | **38** | **345.4Mb** | 20/93 | 26 | **286.0 Mb** | 21/94 |
| **5% Cell-line DNA** | **33** | **274.4Mb** | 15/90 | 25 | **305.8 Mb** | 18/76 |
| **1% Cell-line DNA** | **67** | **2770.5Mb** | 36/22 | 15 | **155.5 Mb** | 2/17 |
| **No Cell-line DNA** | **5** | **130.6Mb** | 0/4 | 7 | **27.8 Mb** | 0/30 |

\* These are 100% by definition as comparing self with self.
\*\*Results from plasma-Seq not available

**Figure 3.**
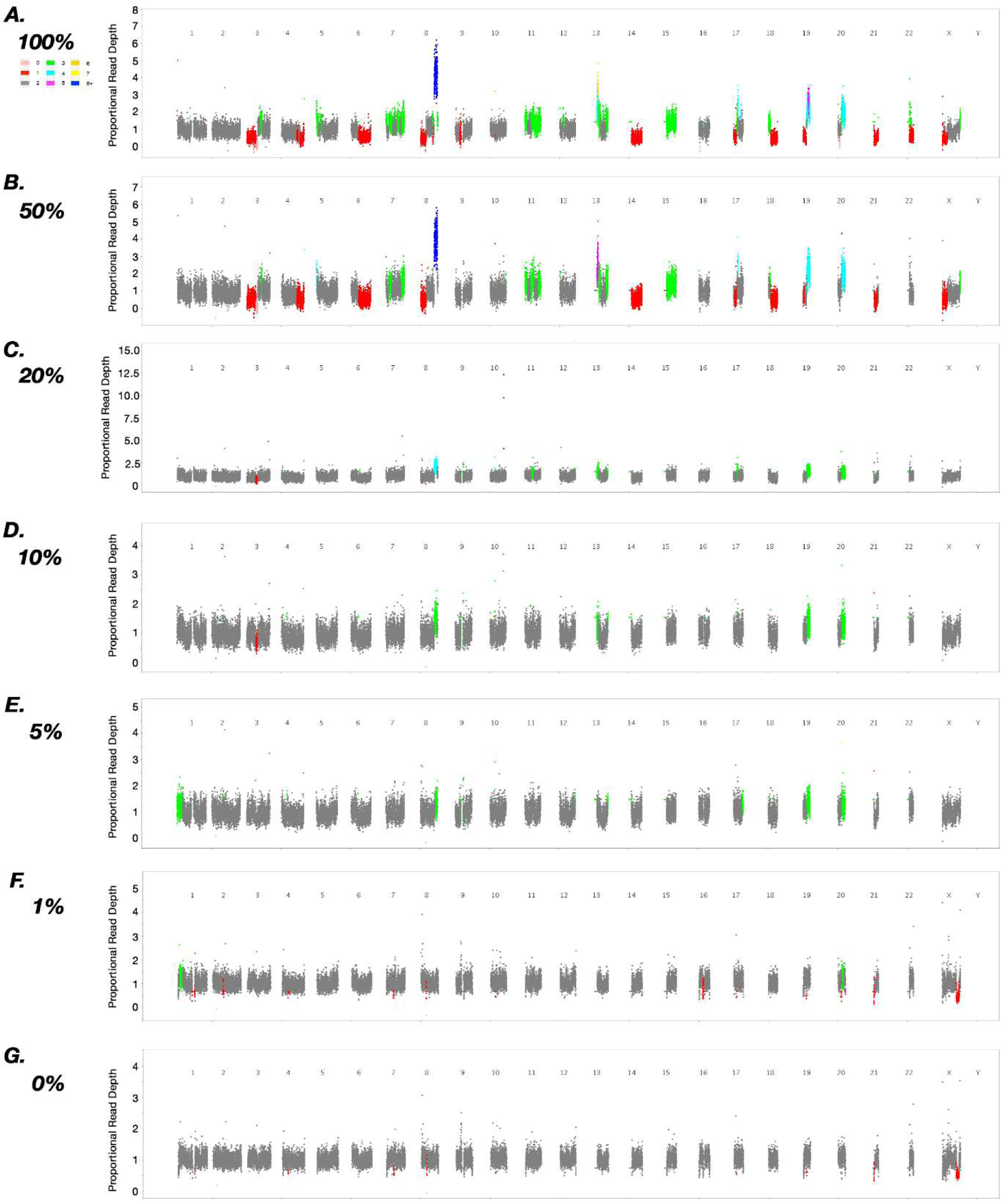
The capacity of the qCNV+sCNA-seq analysis to resolve sCNA across a range of tumour DNA proportion. As shown in the legend, the different colours represent different copy number states. (A) plasma-like sequencing data from a sample that contains 100 % tumour cell-line DNA. The mixtures shown in (B-G) show a range of mixtures containing different proportions of plasma-like tumour cell-line DNA and a plasma-like normal DNA. (B) Mixture containing 50 % cell-line DNA. (C) Mixture containing 20 % cell-line DNA. (D) Mixture containing 10 % cell-line DNA. (E) Mixture containing 5 % cell-line DNA. (F) Mixture containing 1 % cell-line DNA. (G) Mixture containing no tumour cell-line DNA. The sCNA identified in the analysis of the negative control, are likely CNVs or artefacts of the different extraction protocols or sequencing conditions. The use of a ‘matched’ reference sample, which would be collected from same individual and sequenced in parallel would remove the misidentification of CNVs.

Encouragingly some of the clinically significant sCNA identified in the pure cell-line, such as the amplification of 8q, a mutation commonly seen across a range of different cancers [39], and the amplification of 19q a region frequently altered in ovarian cancers [40], were identified in all the mixtures containing more than 5% tumour cfDNA. (Figure. 3A-E), indicating that while a significant proportion of the sCNA seen in the pure cell-line sample were not identified in these samples, there was still enough information to identify biologically relevant mutations in these low ctDNA samples. Furthermore, the 10% mixture revealed that the plasma-Seq analysis captured the copy number profile of the pure cell-line with 20 % sensitivity and 93 % specificity at a single base pair resolution (Table 2). In comparison, the results from the qCNV+sCNA-seq analysis of the 10 % mixture, revealed it reflected the results from the pure cell-line with 21 % sensitivity and 94 % specificity (Table 2), suggesting that in these low purity samples, a single reference sample collected from genomic DNA can perform equally as well as a panel of cfDNA samples collected from healthy individuals.

### A single reference can identify clinically-significant copy numbers in clinical samples

Having demonstrated the potential of our pipeline on mixtures of plasma-like cell-line DNA, we next applied it to characterise publicly available cell free DNA data collected from cancer patients. We re-analysed a cohort of ultra-low coverage, WGS cfDNA samples collected from a group of patients diagnosed with prostate cancer and previously characterised by plasma-Seq [8]. To achieve this, we made use of the same, unmatched blood-derived reference described above. In this case, the reference sample is not sex-matched, as can be observed from the apparent copy number alteration on the X chromosome (Figure 4). Examination of the mutations identified from this cohort revealed sCNA commonly associated with prostate cancer such as the chromosome-scale amplification of the q arm of chromosome 8 (Figure 4A,4D-4F) as well as the deletion of 8p (Figure. 4A, 4D) and 2q (Figure 4A,4G) [41]. These mutations were all evident in the original plasma-Seq analysis of this cohort.

**Figure 4.**
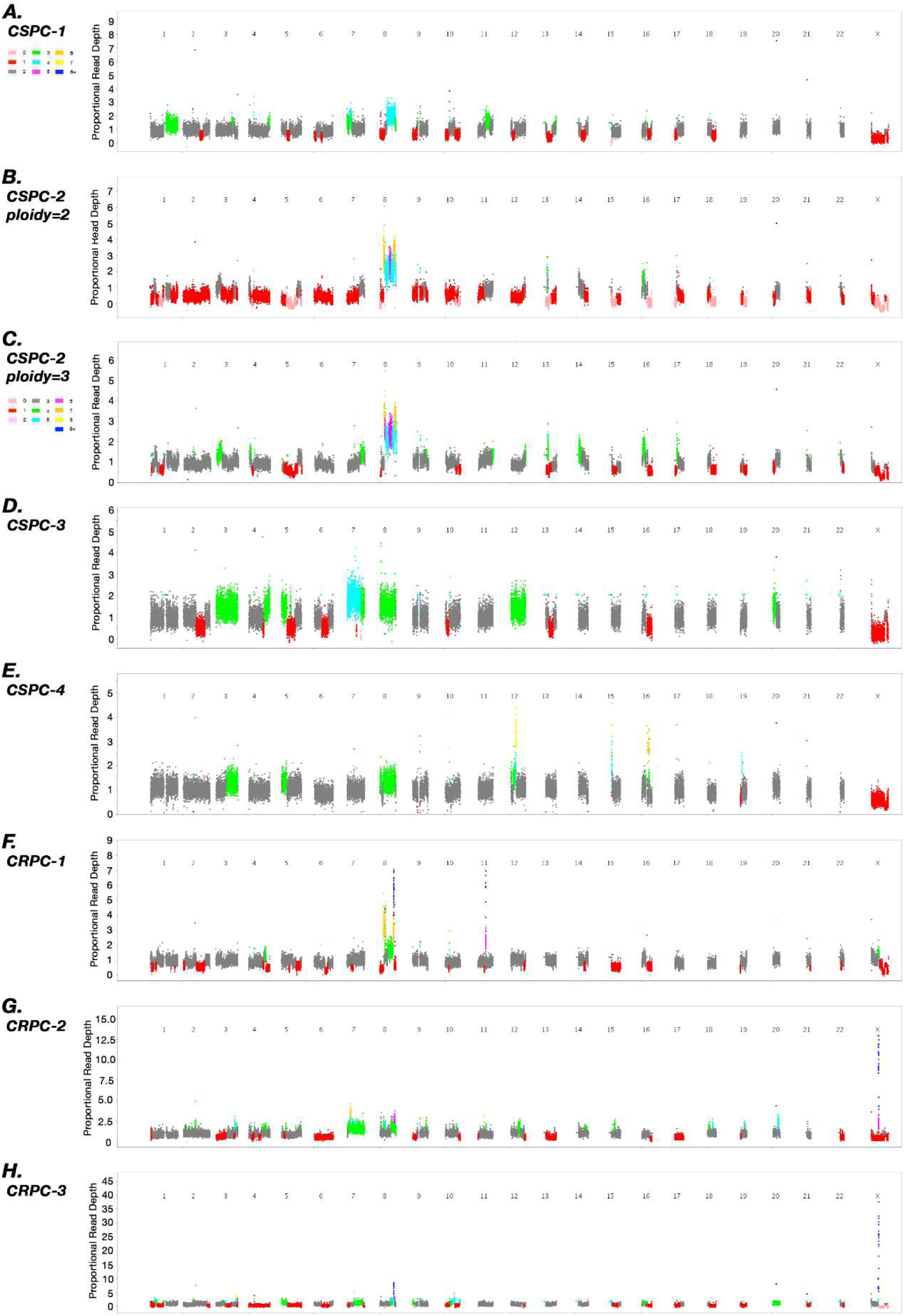
Copy number profile of Castration Sensitive (CS) and Castration Resistant (CR) Prostate Cancers (PC) identified by qCNV+sCNA-seq analysis of cfDNA. The different colours represent different copy number states, with the exception of C, all other images assume these samples were diploid and use the colours shown in legend of A. A-E copy number profile from castration sensitive prostate cancer samples. C copy number profile from CSPC2 that has been reanalysed with increased ploidy. F-H castration resistant prostate cancer samples. The amplification of the Androgen Receptor in chromosome X -an established therapeutic evasive mutation is clearly visible in G and H.

This cohort included samples from both castration sensitive (CS) and castration resistant (CR) forms of the disease. Analysis of the results from two of the patients with CR prostate cancer, revealed a large focal peak of amplification containing the Androgen Receptor on the X chromosome [5] (Figure 4 BC)(Supplementary Figure 2). The amplification of this gene, is an known mechanism of chemo-resistance in the transition of CS to CR prostate cancer [42]. While the remaining CR prostate cancer sample contained prominent focal peaks on chromosome 8 and chromosome 11. Examination of these peaks revealed they contained multiple genes that have been linked to cancer including *CCDN1* as well as genes from the Fibroblast Growth Factor (FGF) family (Supplementary Figure 3). As FGF signalling has been suggested as a novel treatment strategy for CR prostate cancer [43], and the role of *CCDN1* has been well established in cancer [44], this further showcases the potential of our approach. The identification of transformative, chemoresistant copy number mutations from the analysis of cfDNA from the plasma of multiple cancer patients, clearly demonstrates the potential our single reference pipeline to provide clinically significant results from the analysis of ultra-low coverage cfDNA samples.

The identification of recurrent sCNA commonly found in prostate cancer genomes suggests that our single reference method is able to identify clinically significant mutations in real plasma samples from cancer patients. To verify the accuracy of these copy number profiles, the sCNA identified by qCNV+sCNA-seq were compared to the plasma-Seq analysis of the same cohort [8]. This comparison revealed both methodologies were generally able to identify the same broad copy number changes in each of these samples (Supplementary Table 3). One sample, CSPC2 had a low level of overlap with the plasma-Seq copy number profile (Supplementary Table 3), and while the same broad trends were present, examination of the results suggested that the base ploidy had been incorrectly assigned (Figure 4B). Re-analysing this sample as a triploid produced results that were much more consistent with the plasma-Seq analysis (Figure 4C, Supplementary Table 3).

### sCNA-seq captures tumour evolution in clinical exome data

This pipeline was used to analyse two cohorts of matched (tumour, plasma, and blood cell derived DNA), high coverage exome samples. The capacity of the cfDNA to reflect the SNV content of each patient’s disease [17] had been previously described for each of these data-sets, however the copy number profile of these samples had not been determined.

Four samples from the first patient had been collected over a 52-month period. These samples represented the DNA extracted from a FFPE primary breast cancer sample, a metastatic liver lesion collected four years after the primary tumour was excised, a cfDNA sample collected two months after the liver biopsy and sample of the patient’s germline DNA collected from white blood cells at the same time as the plasma sample. The coverage of these samples ranged from 118x to 309x. The combination of these samples made it possible to determine how the copy number profile of the patient’s disease changed over this time, and explore the capacity of exome sequencing to reflect these changes through the analysis of the cfDNA. Characterisation of the primary sample and the metastatic lesion with qCNV+sCNA-seq, both identified the same general trends, including specific scenas such as the focal amplification on chromosome 19 and the deletion of the p arm of chromosome 8 (Figure 5AB). In addition to these events, the metastatic lesion also contained a number of events not present in the primary tumour (Figure 5AB) such as a number of focal amplifications on the p arm of chromosome 11, a result consistent with evolution of the patient’s disease over a four-year time period [37].

**Figure 5.**
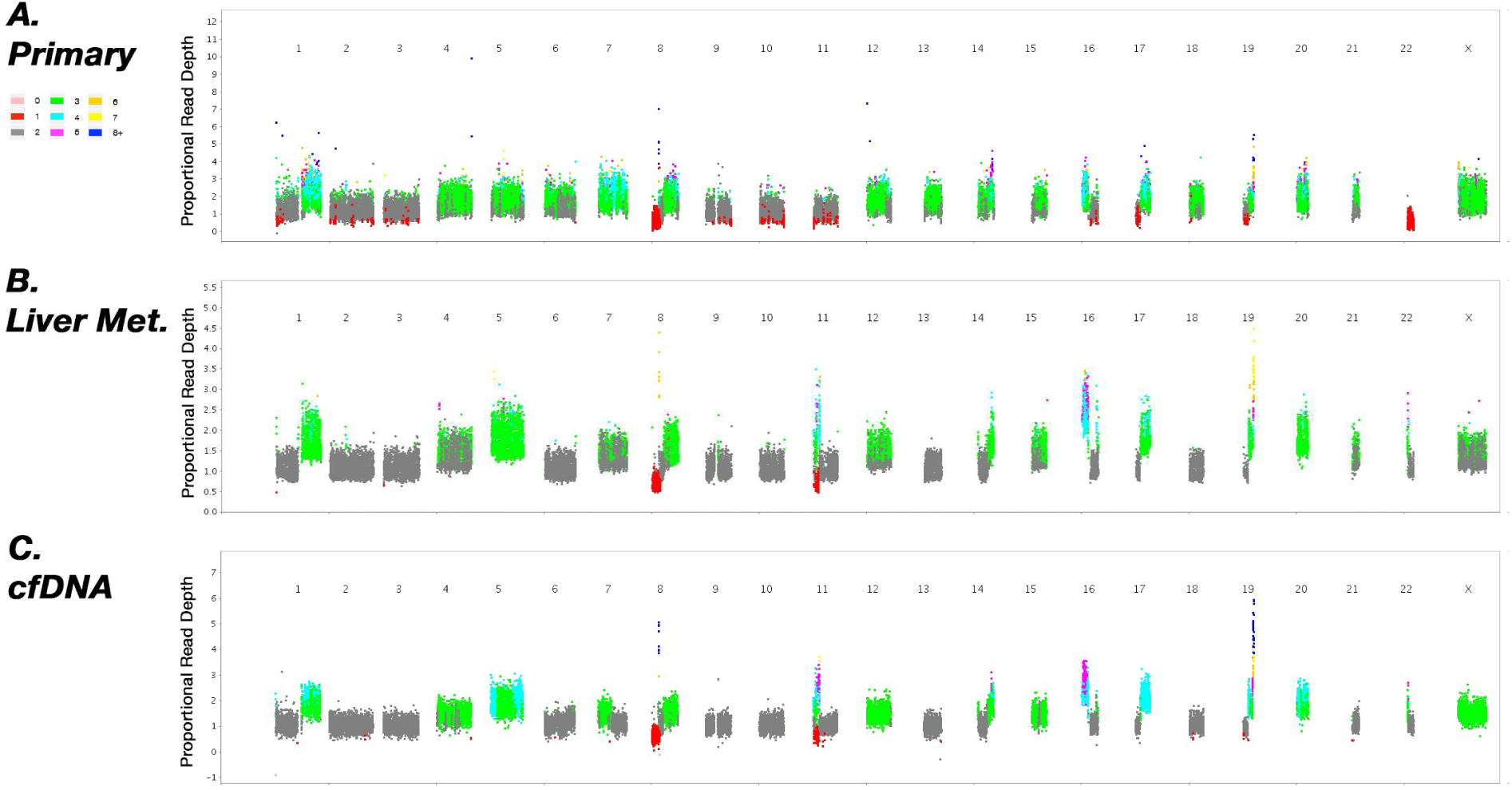
The identification of sCNAs from pairs is dependent on the quality of normal and plasma sample. A. The Primary Breast Cancer vs. matched normal comparison shows a clear copy number profile, despite being undergoing a formalin-fixed, paraffin embedded process. B. The Metastatic Breast Cancer vs matched normal reflects the copy number profile of the primary tumour, but also captures a number of unique of sCNAs, such as the focal amplification on chromosome 11 and the gain of chromosome 22. C. The capacity of the cfDNA sample vs. matched normal to reflect the same broad copy number profile seen in the primary and metastatic samples, clearly demonstrates the clinical utility of our single reference methodology.

Analysis of the results from the cfDNA sample (Figure. 5C) revealed that this analysis was able to recapitulate the same general copy number profile present in the primary and metastatic samples (Figure. 5AB). The sCNAs captured in this analysis included the same current focal amplifications and large-scale changes identified in the primary and metastatic samples, as well as the sCNAs unique to the metastatic lesion. Furthermore, this comparison also identified sCNAs not seen in either solid tumour sample, potentially suggesting the ability of cfDNA to provide a view of the global copy number profile a patient’s disease, hinting at the power of cfDNA for non-invasive monitoring. Together, these samples demonstrate the capacity of qCNV+sCNA-seq to effectively profile a patient’s disease through copy number profiling of solid and cell-free samples, as well as highlighting the potential of a single, matched normal sample to act as an effective reference in real, clinical data.

In contrast, the results from the analysis of the samples from the second patient failed to provide the same level of insight (Supplementary Figure 4). Three samples were collected for sequencing from the second patient, a sample from a primary sarcoma, the cfDNA in the plasma and the patient’s matched normal, germline DNA. Both primary sarcoma, and cfDNA comparisons failed the internal QC check (Supplementary Table 4). Investigating these samples revealed differences in capture array and the resulting sequencing that may have affected the distribution of reads across these libraries (Supplementary Figure 5). These results show that a single reference is capable of identifying clinically significant sCNA from the cell-free DNA, however, the failure of the sarcoma samples indicates the importance of robust quality control measures when profiling sCNAs.

## Discussion

Liquid biopsies have the potential to transform precision medicine. The current best practice for identifying sCNA in cfDNA samples is plasma-Seq [8], a powerful method, designed specifically to identify sCNA from cfDNA samples. While new tools able to identify sCNA from cfDNA samples have recently been developed [45], like plasma-Seq these methods require a reference panel of cfDNA samples taken from a group of healthy individuals. This approach can be difficult to produce for smaller laboratories to source, or prohibitively expensive depending on the sequencing technology used. In this study, we have developed a model which is capable of producing a cfDNA sCNA segmentation using only a single reference, which produces results comparable to an established Z-score based approach. Moreover, we have demonstrated that it is possible to use buffy-coat derived normal, germline DNA as a reference, instead of plasma DNA from healthy reference sample. This provides a more streamlined protocol for sCNA profiling in cell-free DNA in which genomic germline DNA from the buffy coat is sequenced at the same time as the cell free DNA in plasma, and makes it easier to ensure that the reference sample is sequenced using exactly the same procedure as the plasma sample. This pipeline also makes it possible to compare matched samples removing the mischaracterization of CNV as sCNA – something not possible with existing Z-score based approaches.

We have shown that this approach can accurately infer the sCNA profile of cfDNA even using ultra-low coverage (0.06 – 0.29X) sequence data generated using a single Illumina MiSeq run, provided the proportion of tumour derived DNA circulating in plasma is 20% or higher. Some clinically relevant, focal amplification could be observed at lower tumour purity (down to 10%); however, a substantial proportion of copy number changes are missed with ultra-low coverage sequencing of low tumour purity samples, both by our approach as well as plasma-Seq. This represents an achievement as traditional, matched copy number profiling approaches rely on SNV frequencies would be unable to identify mutations at this low level of coverage. While sCNA-Seq is currently unable to deduce the ploidy of a tumour sample apriori, the low level of coverage needed to identify clinically significant mutations in low purity samples, even when ploidy has been mis-assigned, highlights the potential of this approach for rapid sCNA profiling.

One way to address lower tumour purity samples is to use high coverage exome sequencing of plasma DNA. We have shown that our approach applied to a high coverage (309x) sequencing of DNA from plasma coupled with a 201x sequencing of normal DNA in the buffy coat could infer tumour sCNA at high resolution in a sample in which 28% of cell free DNA was tumour derived [17].

Many of the comparisons described here relied on a single unmatched plasma cfDNA reference. This approach identified mutations known to be involved in cancer, however it also has the potential to conflate germline CNV differences with sCNA. A recent estimate suggested that between 4.8 – 9.5% of the genome is affected by CNV [46]. These germ-line differences, begin to influence the signal in cases in which the tumour purity is very low, and/or does not contain significant somatic copy number changes. To avoid the potential for detecting germline CNV we recommend using matched normal DNA as the reference sample.

The detection of therapeutically relevant sCNA from clinical cfDNA samples collected from patients with prostate cancer demonstrates the potential clinical utility of the qCNV+sCNA pipeline. This potential was best exemplified by the identification of the causative, chemo-resistant mutations from the cfDNA of two patients with CR prostate cancer. Even when the base-line ploidy was misassigned in CSPC-2, it was still possible to identify the focal peaks of amplification, more importantly these peaks were still readily identifiable, when this sample was re-analysed as a triploid. The identification of some of the clinically significant sCNA in the low tumour content mixtures, as well as the results from the prostate cancer patients highlights the potential for the qCNV+sCNA-Seq pipeline in cfDNA studies.

## Conclusion

The ability to profile sCNA from a single-sample unlocks the potential of liquid biopsy both to smaller labs, and also for further experimentation using new platforms. By making a version of this approach available on Galaxy, we have also lowered the bioinformatics barriers to using this approach.

## Availability

sCNA-Seq can be downloaded from https://github.com/lachlancoin/scnaseq

qCNV is available at https://sourceforge.net/p/adamajava/wiki/qCnv/

## Acknowledgements

This work was supported by the Queensland Cancer Council with grant number APP1085786. The sequence data from fragmented cfDNA-like samples and the results from the plasma-Seq analysis of these samples was provided by Ellen Heitzer. The authors greatly appreciate this contribution. The authors also appreciate the COSMIC Cancer Cell Line Encyclopaedia for providing the array-based copy number profile of the cell-line, and the use of the raw WGS data from Bulter. et al. The authors wish to acknowledge advice of Minh Duc Cao in the preparation of this manuscript. This research was supported by the use of the NeCTAR Research Cloud, by QCIF and by the University of Queensland’s Research Computing Centre (RCC). The NeCTAR Research Cloud is a collaborative Australian research platform supported by the National Collaborative Research Infrastructure Strategy. The authors also wish to acknowledge the patients who donated tissue and blood samples for the benefit of this research.

## Supplementary Figures

**Supplementary Figure 1.**
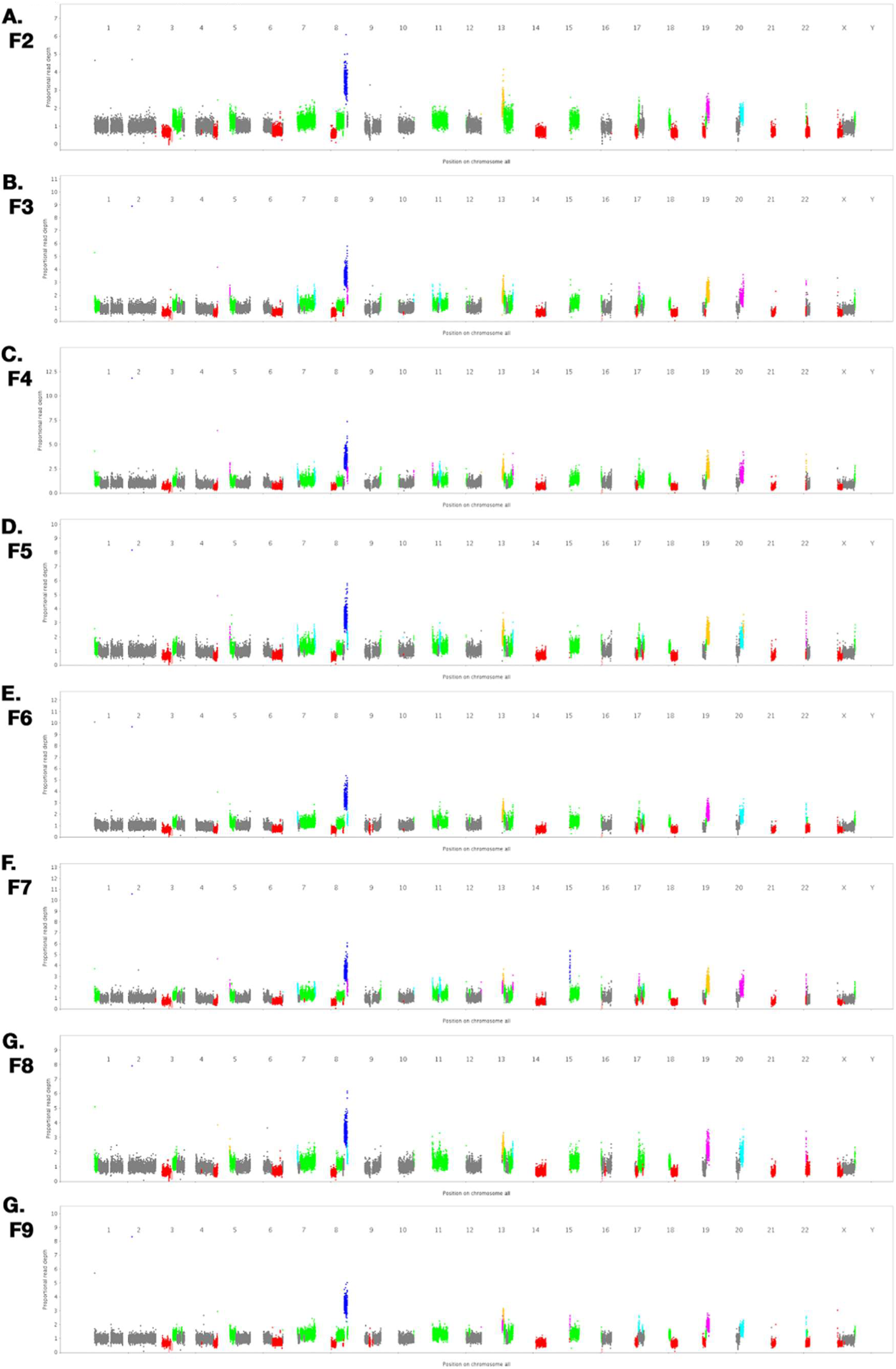
The copy number profile of the pure, plasma like cell-line and each of the normal controls used in the plasma-seq publication.

**Supplementary Figure 2.**
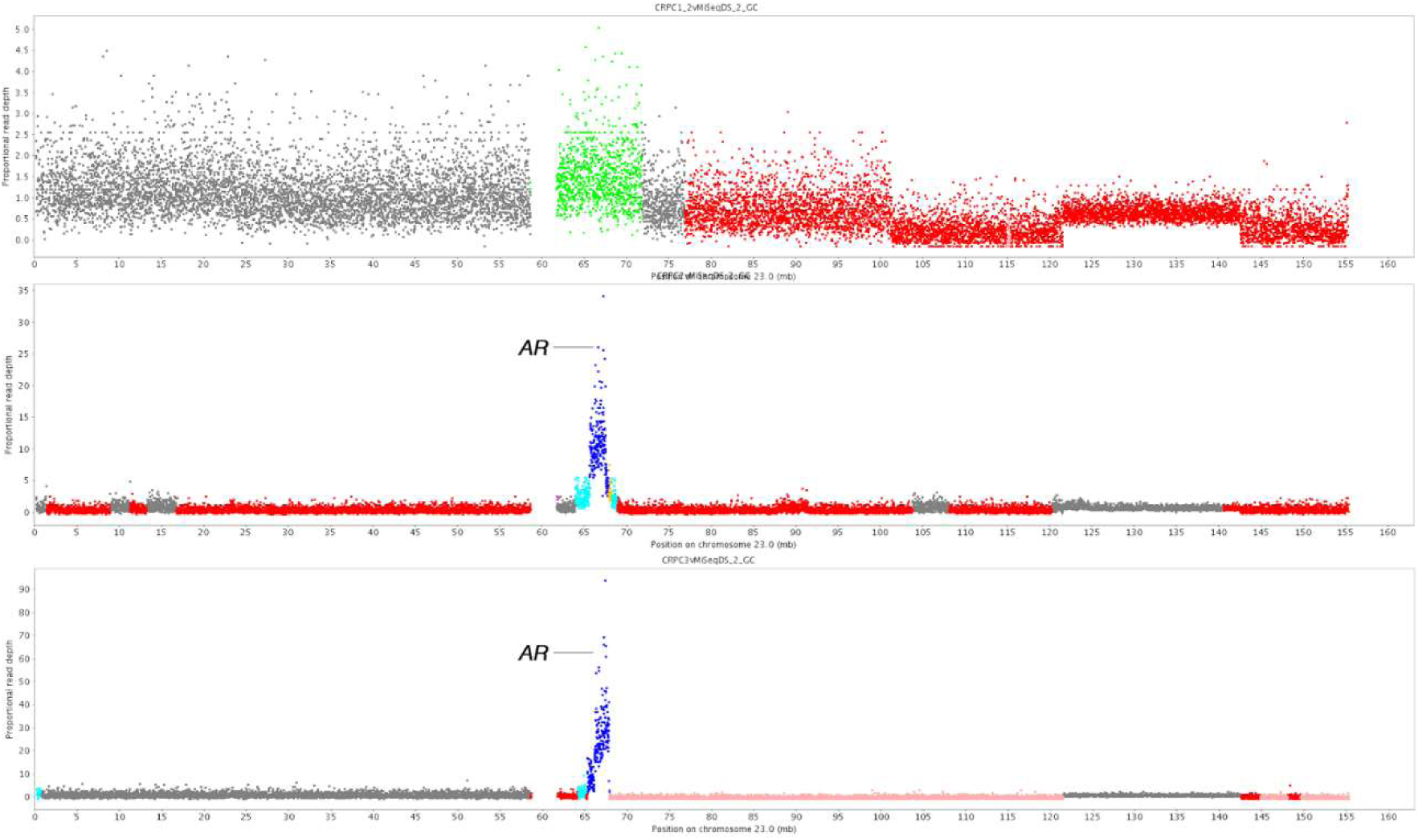
The copy number profile of chromosome X from the analysis of the cfDNA from patients with castration resistant prostate cancer. B and C, clearly show the amplification of a region containing the Androgen Receptor, an established transformative mutation. This clearly shows the potential of qCNV+sCNA-Seq to monitor a patient’s disease non-invasively. A shows the copy number profile from CRPC-1 and demonstrates that there are multiple routes towards castration resistance.

**Supplementary Figure 3.**
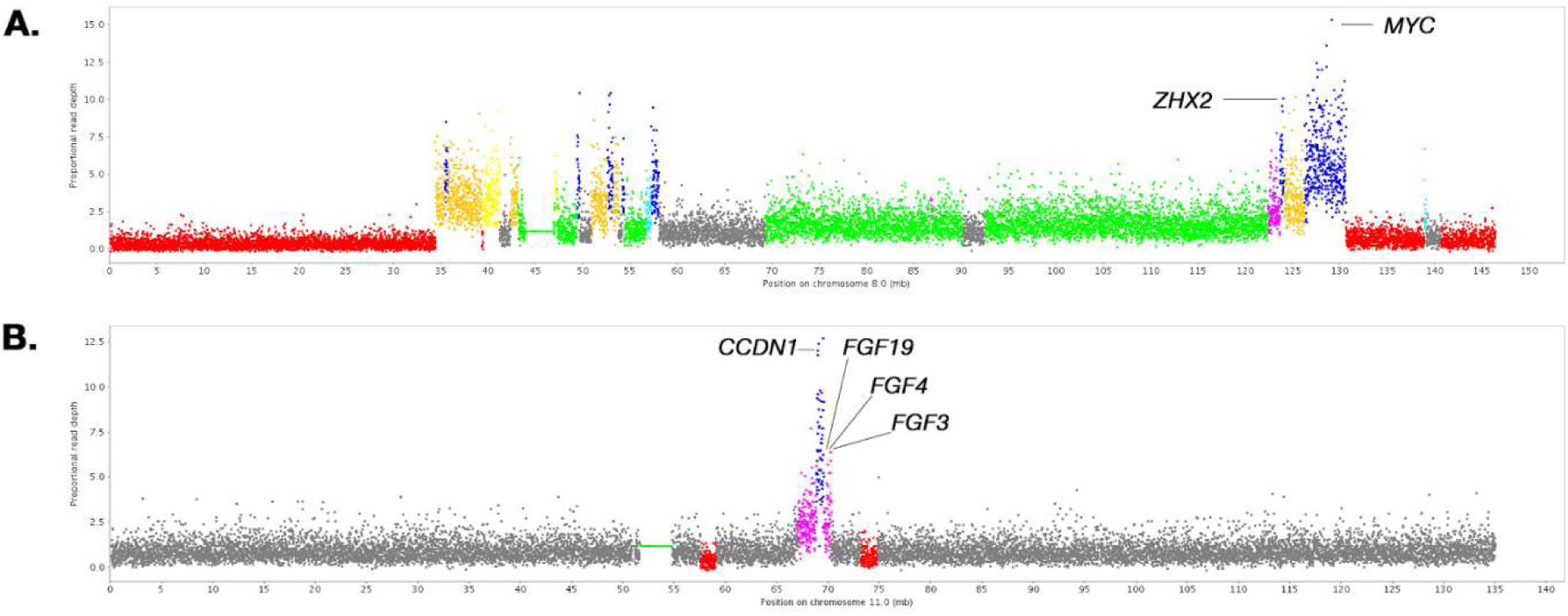
The focal peaks of amplifications from the cfDNA of CRPC-1. Analysis of cfDNA from CRPC1 revealed two prominent peaks of amplification on chromosome 8 (A) and chromosome 11 (B). Examination of these peaks revealed they contained a number of genes strongly associated with cancer, including *MYC*, and *CCDN1*.

**Supplementary Figure 4.**
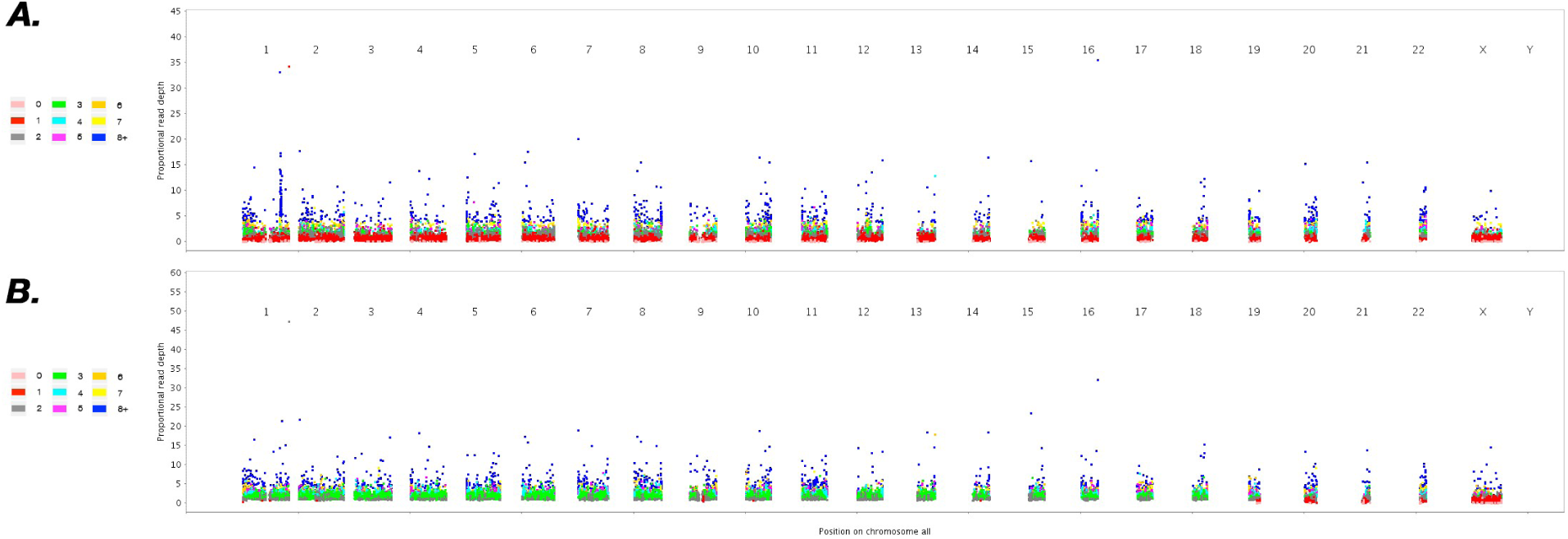
Uninformative copy number profiles from the sarcoma patient. A. Shows the copy number profile from the primary tumour. B shows the copy number profile from the cell-free DNA.

**Supplementary Figure 5.**
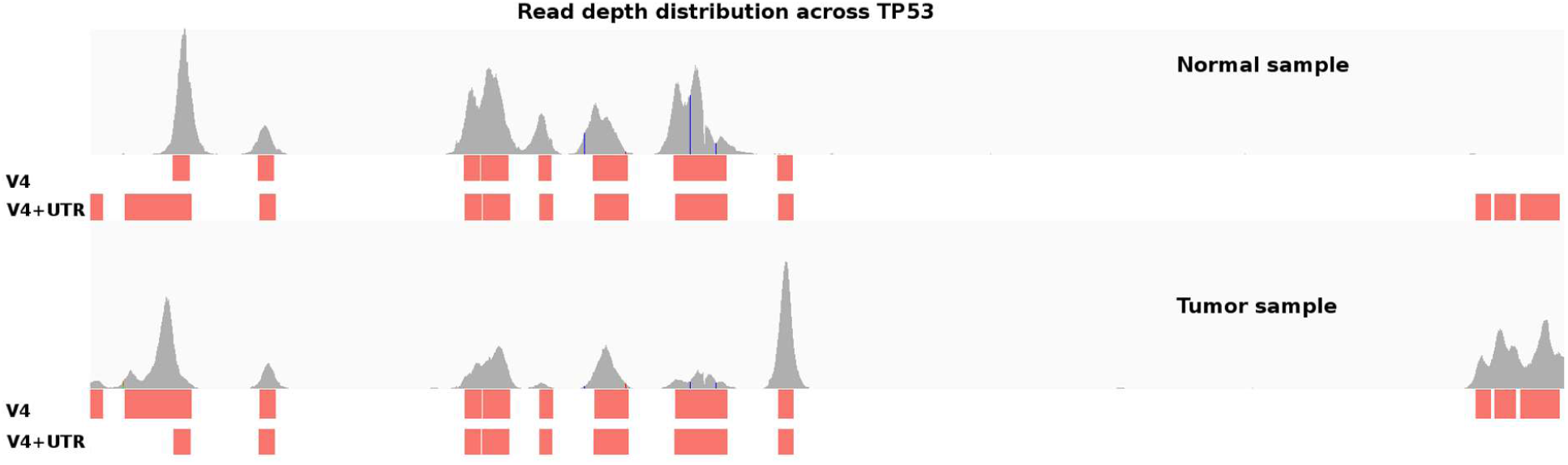
Read depth distribution for the normal and primary tumour sample taken from the second patient in the Butler cohort, compared to the positions of the different probes used in the V4 and V4+UTR exon capture arrays.

## Supplementary Methods

In these experiments, sCNA-Seq was run in distinct stage, or modes. Mode 1 was used to define the purity and ratio of the sample using a larger window size and a small number of mutational states, across multiple iterations. This information was used in the Whole Genome Characterisation Mode (Mode 2) and the High Resolution Regional Survey Mode (Mode 3). Both these modes were run with a single iteration and used a significantly smaller window size as well as a larger number of mutational states.

### Mode 1 -Whole Genome Training

- Window Size = 250 Kb
- Maximum number of potential copy number states = 2 x base ploidy
- Number of iterations = 25
- BetaDownWeight = 10

### Mode 2 -Whole Genome Characterisation

- Window Size = 100 Kb
- Maximum number of potential copy number states = 5 x base ploidy
- Number of iterations = 1
- BetaDownWeight = 10

### Mode 3 -High Resolution, Regional Survey

- Window Size = 5 Kb (variable)
- Maximum number of potential copy number states = 5 x base ploidy
- Number of iterations = 25
- BetaDownWeight = 2
- Specific chromosomal region < 250Mb

## References

1. Hanahan, D. and R.A. Weinberg, The hallmarks of cancer. Cell, 2000. 100(1): p. 57–70.

2. Hanahan, D. and R.A. Weinberg, Hallmarks of cancer: the next generation. Cell, 2011. 144(5): p. 646–74.

3. Slamon, D.J., et al., Human-Breast Cancer - Correlation of Relapse and Survival with Amplification of the Her-2 Neu Oncogene. Science, 1987. 235(4785): p. 177–182.

4. Carter, P., et al., Humanization of an anti-p185HER2 antibody for human cancer therapy. Proc Natl Acad Sci U S A, 1992. 89(10): p. 4285–9.

5. Visakorpi, T., et al., In-Vivo Amplification of the Androgen Receptor Gene and Progression of Human Prostate-Cancer. Nature Genetics, 1995. 9(4): p. 401–406.

6. Feldman, B.J. and D. Feldman, The development of androgen-independent prostate cancer. Nat Rev Cancer, 2001. 1(1): p. 34–45.

7. Chan, K.C., et al., Cancer genome scanning in plasma: detection of tumor-associated copy number aberrations, single-nucleotide variants, and tumoral heterogeneity by massively parallel sequencing. Clin Chem, 2013. 59(1): p. 211–24.

8. Heitzer, E., et al., Tumor-associated copy number changes in the circulation of patients with prostate cancer identified through whole-genome sequencing. Genome Med, 2013. 5(4): p. 30.

9. Heitzer, E., P. Ulz, and J.B. Geigl, Circulating tumor DNA as a liquid biopsy for cancer. Clin Chem, 2015. 61(1): p. 112–23.

10. Heitzer, E., et al., Non-invasive detection of genome-wide somatic copy number alterations by liquid biopsies. Mol Oncol, 2016. 10(3): p. 494–502.

11. Thierry, A.R., et al., Clinical validation of the detection of KRAS and BRAF mutations from circulating tumor DNA. Nature Medicine, 2014. 20(4): p. 430-+.

12. Karapetis, C.S., et al., K-ras mutations and benefit from cetuximab in advanced colorectal cancer. New England Journal of Medicine, 2008. 359(17): p. 1757–1765.

13. Misale, S., et al., Emergence of KRAS mutations and acquired resistance to anti-EGFR therapy in colorectal cancer. Nature, 2012. 486(7404): p. 532–6.

14. Mohan, S., et al., Changes in colorectal carcinoma genomes under anti-EGFR therapy identified by whole-genome plasma DNA sequencing. PLoS Genet, 2014. 10(3): p. e1004271.

15. Forshew, T., et al., Noninvasive identification and monitoring of cancer mutations by targeted deep sequencing of plasma DNA. Sci Transl Med, 2012. 4(136): p. 136ra68.

16. Bettegowda, C., et al., Detection of circulating tumor DNA in early- and late-stage human malignancies. Sci Transl Med, 2014. 6(224): p. 224ra24.

17. Butler, T.M., et al., Exome Sequencing of Cell-Free DNA from Metastatic Cancer Patients Identifies Clinically Actionable Mutations Distinct from Primary Disease. PLoS One, 2015. 10(8): p. e0136407.

18. Norwitz, E.R. and B. Levy, Noninvasive prenatal testing: the future is now. Rev Obstet Gynecol, 2013. 6(2): p. 48–62.

19. Pescia, G., et al., Cell-free DNA testing of an extended range of chromosomal anomalies: clinical experience with 6,388 consecutive cases. Genet Med, 2016.

20. Heidary, M., et al., The dynamic range of circulating tumor DNA in metastatic breast cancer. Breast Cancer Res, 2014. 16(4): p. 421.

21. Ulz, P., et al., Whole-genome plasma sequencing reveals focal amplifications as a driving force in metastatic prostate cancer. Nat Commun, 2016. 7: p. 12008.

22. Heitzer, E., et al., Circulating tumor cells and DNA as liquid biopsies. Genome Med, 2013. 5(8): p. 73.

23. Giardine, B., et al., Galaxy: a platform for interactive large-scale genome analysis. Genome Res, 2005. 15(10): p. 1451–5.

24. Lappalainen, I., et al., The European Genome-phenome Archive of human data consented for biomedical research. Nat Genet, 2015. 47(7): p. 692–5.

25. Leinonen, R., et al., The European Nucleotide Archive. Nucleic Acids Res, 2011. 39(Database issue): p. D28–31.

26. Forbes, S.A., et al., COSMIC: exploring the world’s knowledge of somatic mutations in human cancer. Nucleic Acids Research, 2015. 43(D1): p. D805–D811.

27. Greenman, C.D., et al., PICNIC: an algorithm to predict absolute allelic copy number variation with microarray cancer data. Biostatistics, 2010. 11(1): p. 164–75.

28. Wickham, H., ggplot2: Elegant Graphics for Data Analysis. Ggplot2: Elegant Graphics for Data Analysis, 2009: p. 1–212.

29. Li, H., et al., The Sequence Alignment/Map format and SAMtools. Bioinformatics, 2009. 25(16): p. 2078–9.

30. Li, H., Aligning sequence reads, clone sequences and assembly contigs with BWA-MEM. arXiv, 2013. 1303.3997v1([q-bio.GN).

31. Picard. 2016, Broad Institute. p. Picard: A set of command line tools (in Java) for manipulating high-throughput sequencing (HTS) data and formats such as SAM/BAM/CRAM and VCF.

32. AdamaJava. https://sourceforge.net/projects/adamajava/ [cited 2017 06-30].

33. Bellos, E., M.R. Johnson, and L.J.M. Coin, cnvHiTSeq: integrative models for highresolution copy number variation detection and genotyping using population sequencing data. Genome Biology, 2012. 13(12).

34. Belic, J., et al., Rapid Identification of Plasma DNA Samples with Increased ctDNA Levels by a Modified FAST-SeqS Approach. Clin Chem, 2015. 61(6): p. 838–49.

35. Kawai, K., et al., Comprehensive karyotyping of the HT-29 colon adenocarcinoma cell line. Genes Chromosomes Cancer, 2002. 34(1): p. 1–8.

36. Corzo, C., et al., RxFISH karyotype and MYC amplification in the HT-29 colon adenocarcinoma cell line. Genes Chromosomes Cancer, 2003. 36(4): p. 425–6.

37. Ulz, P., M. Auer, and E. Heitzer, Detection of Circulating Tumor DNA in the Blood of Cancer Patients: An Important Tool in Cancer Chemoprevention. Methods Mol Biol, 2016. 1379: p. 45–68.

38. Ulz, P., et al., Inferring expressed genes by whole-genome sequencing of plasma DNA. Nat Genet, 2016. 48(10): p. 1273–8.

39. Zack, T.I., et al., Pan-cancer patterns of somatic copy number alteration. Nat Genet, 2013. 45(10): p. 1134–40.

40. Bayani, J., et al., Genomic instability and copy-number heterogeneity of chromosome 19q, including the kallikrein locus, in ovarian carcinomas. Mol Oncol, 2011. 5(1): p. 48–60.

41. Williams, J.L., P.A. Greer, and J.A. Squire, Recurrent copy number alterations in prostate cancer: an in silico meta-analysis of publicly available genomic data. Cancer Genet, 2014. 207(10-12): p. 474–88.

42. Koivisto, P., et al., Androgen receptor gene amplification: a possible molecular mechanism for androgen deprivation therapy failure in prostate cancer. Cancer Res, 1997. 57(2): p. 314–9.

43. Gowardhan, B., et al., Evaluation of the fibroblast growth factor system as a potential target for therapy in human prostate cancer. British Journal of Cancer, 2005. 92(2): p. 320–327.

44. Motokura, T., et al., A novel cyclin encoded by a bcl1-linked candidate oncogene. Nature, 1991. 350(6318): p. 512–5.

45. Adalsteinsson, V.A., et al., Scalable whole-exome sequencing of cell-free DNA reveals high concordance with metastatic tumors. Nature Communications, 2017. 8(1): p. 1324.

46. Zarrei, M., et al., A copy number variation map of the human genome. Nat Rev Genet, 2015. 16(3): p. 172–83.

